# Agreement Test Between The Six Minutes Walking Test And Four Meter Gait Speed

**DOI:** 10.1101/2020.11.20.391193

**Authors:** Nury Nusdwinuringtyas, Tresia Fransiska, Peggy Sunarjo, Kevin Triangto, Sopiyudin Dahlan

## Abstract

**Background/Objective:** In the field of Physical Medicine and Rehabilitation, it is essential to measure individual functional capacity, which could be evaluated through walking tests. Aside from the commonly used six minutes walking test (6MWT), four meter gait speed (4MGS) are widely used for its practicality. This study aimed to assess the agreement between 4 MGS and the 6MWT in Indonesian healthy adults.

**Methods:** This agreement analysis study had recruited 61 healthy and sedentary Indonesians aged 18 until 50 years old, and they were instructed to perform three tests, namely 6MWT and 4MGS with six meters and eight meters track. These gait speed were then compared to assess validity.

**Results:** Mean gait speed results for males in 6MWT is 1.602 m/s, whereas 4MGS in six meter track is 2.114 m/s and similarly 2.108 m/s in the eight meter track. Females on the other hand, achieved 1.462 m/s for 6MWT, 1.908 m/s and 1.986 m/s for 4MGS in six and eight meter simultaneously. Bland Altman Agreement test between the 6MWT and 4MGS shows scatter dots with close limit of agreement, thus showing a good agreement between the 6MWT and 4 MGS with both tracks.

**Discussion:** Both track length of 4 MGS were in a good agreement with 6MWT for functional capacity assessment.

**Conclusions:** In response to the COVID-19 pandemic era, shorter track of 4MGS (six meters) can be feasibly utilized. It is evident that shorter duration and track will boost the tests practicality in assessing functional capacity for both inpatient and outpatient settings.

## Introduction

The six minute walk test (6MWT) is a functional capacity test which is widely used, as walking depicts a general activity in daily living.^1,2^ This test is also classified as a submaximal exercise test with walking distance as its outcome.^3^ Originally, the American Thoracic Society (ATS) recommended a 30 meter^1^ continuous track to perform the 6MWT, while more recent studies by Nury revealed the 6MWT validity when conducted in a shorter 15 meter track.^3,4^ It was shown previously that track length would correlate to distance travelled, hence it became an issue to be addressed.^5^

Aside from distance based outcome, other walk tests have gait speed as its outcome, stating the fact that gait speed highly correlates to an individual’s functional capacity. Therefore, gait speed has also been considered as the sixth vital sign.^6,7^ Although it is easy to assess, gait speed depicts functional abilities of various organ systems, such as sensory, motoric, perceptual, cognitive, musculoskeletal, mental status, and energy generation. In particular, energy generation involves both cardiorespiratory and metabolic system, which then highly correlates to an individual’s activity level. Gait speed would then decrease if one or more of the aforementioned functions are disturbed. Hence, gait speed has also become several predictors including disability, cognitive disturbance, fall, and mortality.^8^ Over the years, one of the most commonly utilized gait speed outcome test is the four meter gait speed test (4MGS)^9^, which is generally performed in a six or eight meter straight walking track.

In the corona virus disease 2019 (COVID-19) pandemic era, it is well known that cardiopulmonary system took the greatest toll, and hence endurance examination is essential to be performed.^10^ The attempts to stem nosocomial COVID-19 transmission begins from setting specified zones in the hospital area, separating those with high probability of infection, and those who are proven to be free of infection.^11,12^ This zonation has greatly impacted the limited space that could be utilized for walking test purposes.^12^

Although the 4MGS could then be seen fitting to be applied during pandemic being a faster and simpler alternative to 6MWT, the agreement have not yet been obtained to the authors’ knowledge. Therefore in the novelty of obtaining a shorter track and a more feasible examination in the pandemic era, this study is aimed to assess the validity and reliability of performing 4MGS in two different track lengths, comparing it to the 6MWT in 15 meter track as a standard, to see which track length for 4MGS best describes the gait speed obtained in 6MWT.

## Methods

This is a cross sectional descriptive study that attempted to obtain the mean gait speed of healthy Mongoloid race Indonesian adults from three walking tests, namely 6MWT in 15 meter track, 4MGS from both six and eight meter track. Analytic study was also performed to assess agreement of gait speed from 6MWT and 4MGS in six and eight meter track.

The target population is healthy adults of 18-50 years old with parents who are both Indonesian. The sample population was then gathered from university students, hospital workers, and the community, all recruited through consecutive sampling.

Sample size calculation will be done by utilizing one sample two sided mean size calculation method:

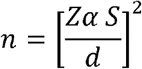

Za = 2.58 (confidence of 99%) and standard deviation (s) of 0.2 m/s (based on a study by Bohannon that comfortable gait speed for both genders age 20-50 years old ranges between 0.094 – 0.229 m/s, thus obtaining the highest value rounded up to 0.2 m/s); and absolute accuracy expected (d) of 0.1 m/s. Acknowledging the cross sectional nature of this study, no drop out was considered. This calculation then yielded a sample size of n = {2.58 × 0.2/(0.1)}^2^ = 26.6 = 27 (rounded up).

This study then attempted to present each gender separately, but the gender classification will not be compared for agreement. Therefore based on the sample size calculation, this study gathered double of the initial calculation accounting for each genders, summed up into 60 subjects. Sample size calculation for both genders are considered the same due to identical standard deviation (s) and absolute accuracy (d) values.

This study was performed in the Medical Rehabilitation Department of the National Referral Hospital of Cipto Mangunkusumo, Jakarta, Indonesia on 19^th^ August 2016 until 9^th^ March 2017. Each of the subjects were instructed to perform 6MWT^3,4,13^ first before proceeding to 4MGS in six meters track, and eight meters track consecutively. These 6MWT test was performed once for every subject, and an adaptation walk test was done right before recording the 4MGS tests. Finally, the 4MGS tests were each performed twice and recorded by two independent recorders, all of these values were then gathered to obtain the mean gait speed.

Univariate data analysis were initially done to observe the categoric descriptive data such as gender, followed by continuous values of age, body weight, height, and gait speed categorized for each gender. There were no statistical test that was performed in gait speed as they were descriptive data. The Bland Altman analysis were finally done in 4MGS values for both six and eight meter track to see each of their agreement to 6MWT.

## Results

Baseline subject descriptive data was obtained and presented in Table 1. The study obtained 61 subjects, accounting for 28 males and 33 females. Mean age for males were 26.25 (±4.55) years, while females were 27.54 (±4.91) years. As for mean height, males were 171.37 (±6.33) cm, which were subsequently taller as compared to females, with mean height 157.54 (±5.52) cm. Body mass index for the male subjects were 21.41 (±1.83) kg/m^2^, while females 21.65 (±1.82) kg/m^2^. Functional capacity of lung were also recorded and presented, as it is one of the inclusion criteria of healthy adults. This study obtained FEV_1_% of both genders were above 90%.

**Table 1.** Descriptive Subject Characteristics based on Anthropometric and Respiratory Values (male = 28; female= 33)

| Variabel | Mean | SD | P<br>value* |
| --- | --- | --- | --- |
| Age (Years) |  |  |  |
| – Male | 26.25 | 4.55 | 0.008 |
| – Female | 27.54 | 4.91 |  |
| **Height (cm)** |  |  |  |
| – Male | 171.37 | 6.33 |  |
| – Female | 157.04 | 5.52 | 0.554 |
| **Weight (kg)** |  |  |  |
| – Male | 67.35 | 7.49 |  |
| – Female | 53.56 | 6.76 | 0.467 |
| **Body Mass Index (kg/m<sup>2</sup>)** |  |  |  |
| – Male | 21.41 | 1.83 |  |
| – Female | 21.65 | 1.82 | 0.667 |
| **FEV 1% prediction** |  |  |  |
| – Male | 102.09 | 10.72 | 0.241 |
| – Female | 99.55 | 13.33 |  |
| **FVC% prediction** |  |  |  |
| – Male | 94.39 | 9.88 | 0.241 |
| – Female | 96.16 | 13.42 |  |
| **FEV 1%** |  |  |  |
| – Male | 94.03 | 9.88 | 0.710 |
| – Female | 96.72 | 13.35 | 0.536 |

Values obtained from the first test, namely 6MWT, were presented in Table 2. Distance travelled was presented in meter, and speed was presented in m/s. Males have achieved 598.90 (±73.51) meter distance travelled and females were 520.42 (±51.00) meter, these were compared with independent t-test to gain a *p*-value of <0.001. Similar comparison (*p* = 0.99) was also obtained in gait speed for 6MWT, which was 1.602 (±0.375) m/s for males and 1.462 (±0.139) m/s for females.

**Table 2.**
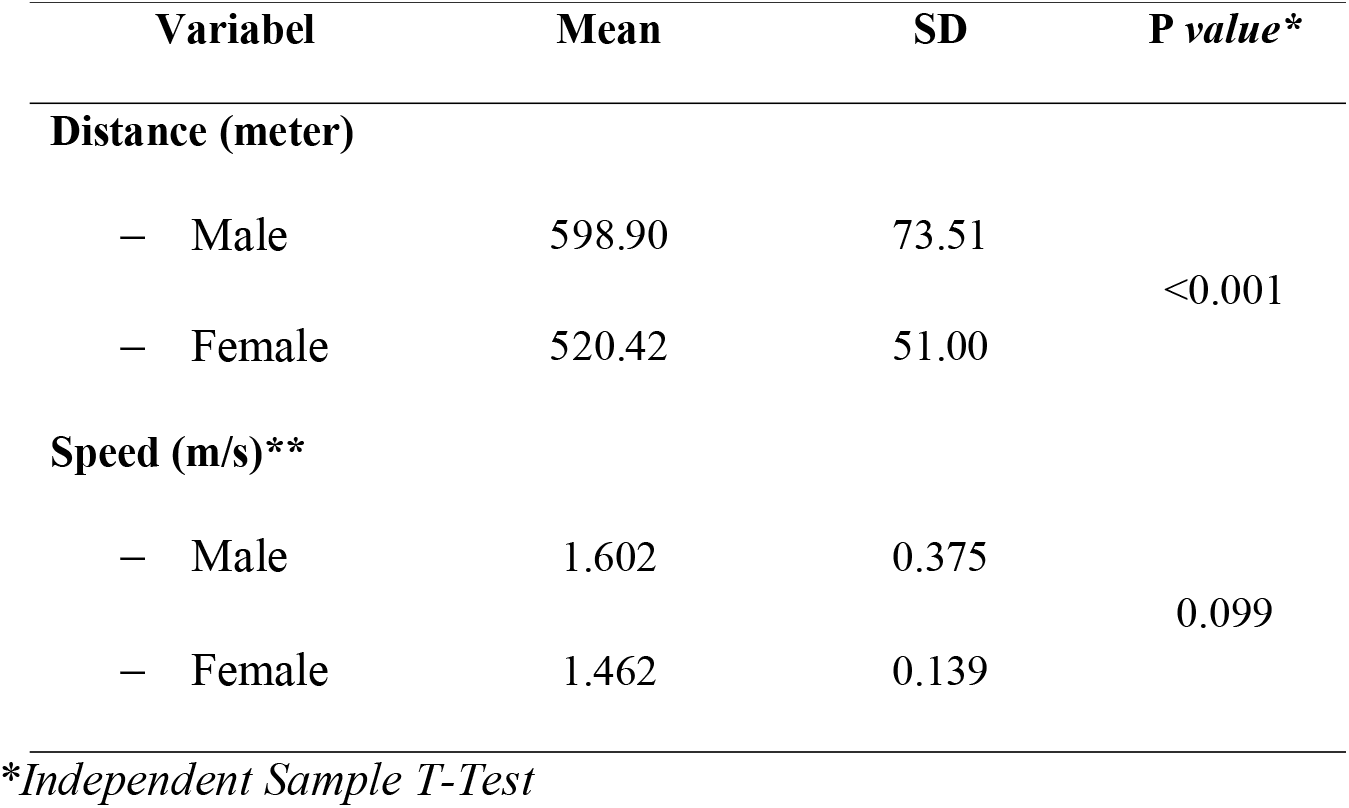
Comparison of Six Minute Walking Test Distance and Speed Values between Gender (male = 28; female= 33)

The 4MGS test values were all presented in Table 3, being categorized for each gender and also track lengths of six and eight meter. Gait speed that was obtained in six meter track for males were 2.114 (±0.309) m/s, as for females were 1.908 (±0.227) m/s. While for 4MGS being performed in eight meter track, males achieved 2.108 (±0.172) m/s, and females 1.986 (±0.339) m/s. All of these tests were compared and were seen to be similar in between gender (p > 0.05), obtaining p value of 0.609 and 0.088 for six and eight meters accordingly.

**Table 3.**
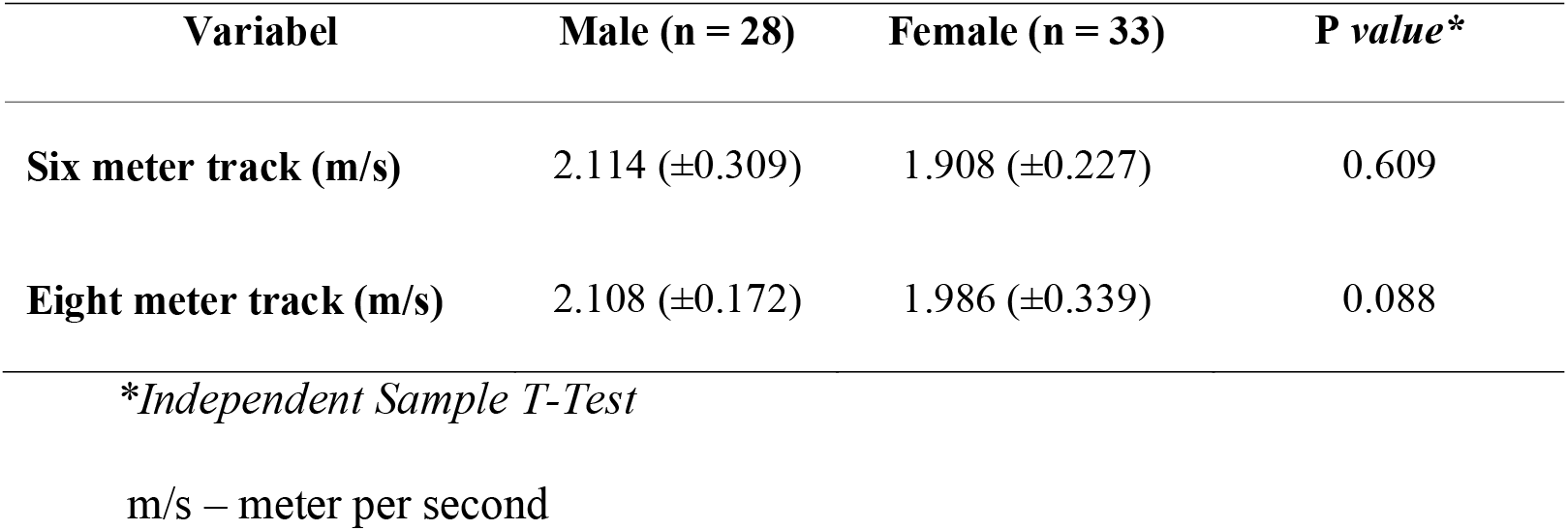
Comparison of 4 Meter Gait Speed on Six and Eight Meter Track between Gender.

Two graphs were also presented to review the gait speed agreement between 6MWT and 4MGS. Graph 1 displayed the results of a Bland Altman gait speed scatterplot for 6MWT and 4MGS in a six meter track. It appeared that the scatter are all within the limit of agreement, showing a valid Bland Altman agreement test. Accordingly, Graph 2 plotted gait speed for 6MWT and 4MGS in <u>eight meter track</u>, with similar results of all the scatter placed within the limit of agreement. Thus for both track lengths, 4MGS are seen to achieve good agreement with 6MWT.

## Discussion

This study has gathered 28 male subjects out of the required 30 subjects, and for female, a total of 33 subjects were recruited out of the required 30 subjects. Overall, the study had met its sample size calculation, as each arm should have 27 subjects minimum.

This study is an extended study from the original 6MWT study with subjects of age 18-50 years, which was translated into the sample age range of this study. Most subjects were under the group of third decade, similar to prior 6MWT studies which obtained the mean age of males 26.78 years while females were 21.92 years old. Male subjects in the study was also seen to be taller, but there was no significant differences between body mass index between the gender. Another gender difference could be seen in 6MWT distance achieved, as males significantly travelled further in comparison to females. This difference depicts gender variability in energy metabolism being more efficient in males, however was not further analysed in this study. However it was seen that gait speed were similar between the gender, and also between 6MWT and 4MGS tests. All these illustrates how gait speed may not correlate directly to distance travelled by 6MWT, but it is enough to reveal the achievement of steady state speed in the tests, showing an overall good agreement.

Gait speed was described as a distance that could be travelled in a specified unit of time, with meter/second or miles/hour being the two most commonly used units. The magnitude will then be greatly affected by anthropometry values such as height or weight, and therefore it should be specified for each race.^14^ Anthropometry will also affect gait speed through step length variability, as they correlate highly to body height.^15^ Additionally, anthropometry will also affect stride length, ultimately resulting in a slower gait in females. This study had shown a tendency of slower female gait speed accounting for taller body height in males.

Gait speed portrays a general condition and is able to predict several clinical outcomes.^6^ It is important to see that differences of 0.1 m/s will have a significant impact to survival rate. Therefore, accuracy of gait speed recording must be given attention, especially for shorter tracks.

Aside from being affected by functional capacity, gait speed also correlates highly to several functional and psychological variables such as age, BMI, FEV1, FVC, residual volume, 6MWT value, total energy expenditure, sedentary lifestyle (<2 METs), number of steps per day, fear of dyspnea and depression.^16^ Gait speed can then be considered as an examination that illustrates a global health condition of an individual, along with multi systemic effects of disease severity, without focusing only on respiratory system.

The 6MWT has been recommended for evaluation of respiratory patients by the ATS.^1^ Additionally, 6MWT is also utilized for other disorders aside from respiratory, but there is a need of an alternative test which are faster and more feasible to be done, owing to a shorter track requirement. Gait speed as mentioned before is an important indicator of functional capacity, and as proven by this study was feasible to be done in both six and eight meter track. It is essential to note the additional distance to the recorded four meter, gives an opportunity for both acceleration and deceleration, in which these are not included in the gait speed measurement. The six meter track was adapted from the National Institutes of Health, providing a one meter acceleration and deceleration, while the eight meter track looked up to National Institute of Ageing (2013) which utilized two meters at each end.^17,18^

Graph 1 effectively portrayed that there were no outliers, putting the fact that 4MGS in a six meter track has a good agreement with 6MWT, as it is within 1.96 standard deviation difference. Similarly in Graph 2, there were no outliers in the eight meter track, and thus also showing good agreement with 6MWT.

Another novelty that is pointed out in this study is the recruitment of specific Mongoloid race of the Indonesian population. There were no biases identified in this recruitment, as inclusion criteria was designed to only include native Indonesian to achieve a higher accuracy of Mongoloid anthropometric values. Gait speed being the final measurements here, is clearly affected by step and stride length, which in turn is affected by anthropometric height measurements.

Essentially as each race varies in the body height, therefore the authors suggest further studies should address walking test values for other race. It is then expected that the results of this study would influence other countries to also set baseline values to ease up practical use of this tool, especially during this pandemic era and its implication for telemedicine purposes.^19^

## Conclusion

Gait speed obtained from 4MGS were seen to be in agreement with the speed of 6MWT performance. Additionally the 4MGS has a shorter test duration and requires a much shorter track as compared to 6MWT. Both of these qualities are valuable during the pandemic era, as the 4MGS would be highly applicable in the telemedicine setting. This study had also shown that both 4MGS track, namely the six and eight meter, have achieved good agreement to the 6MWT.

---

**Grafik 1.**
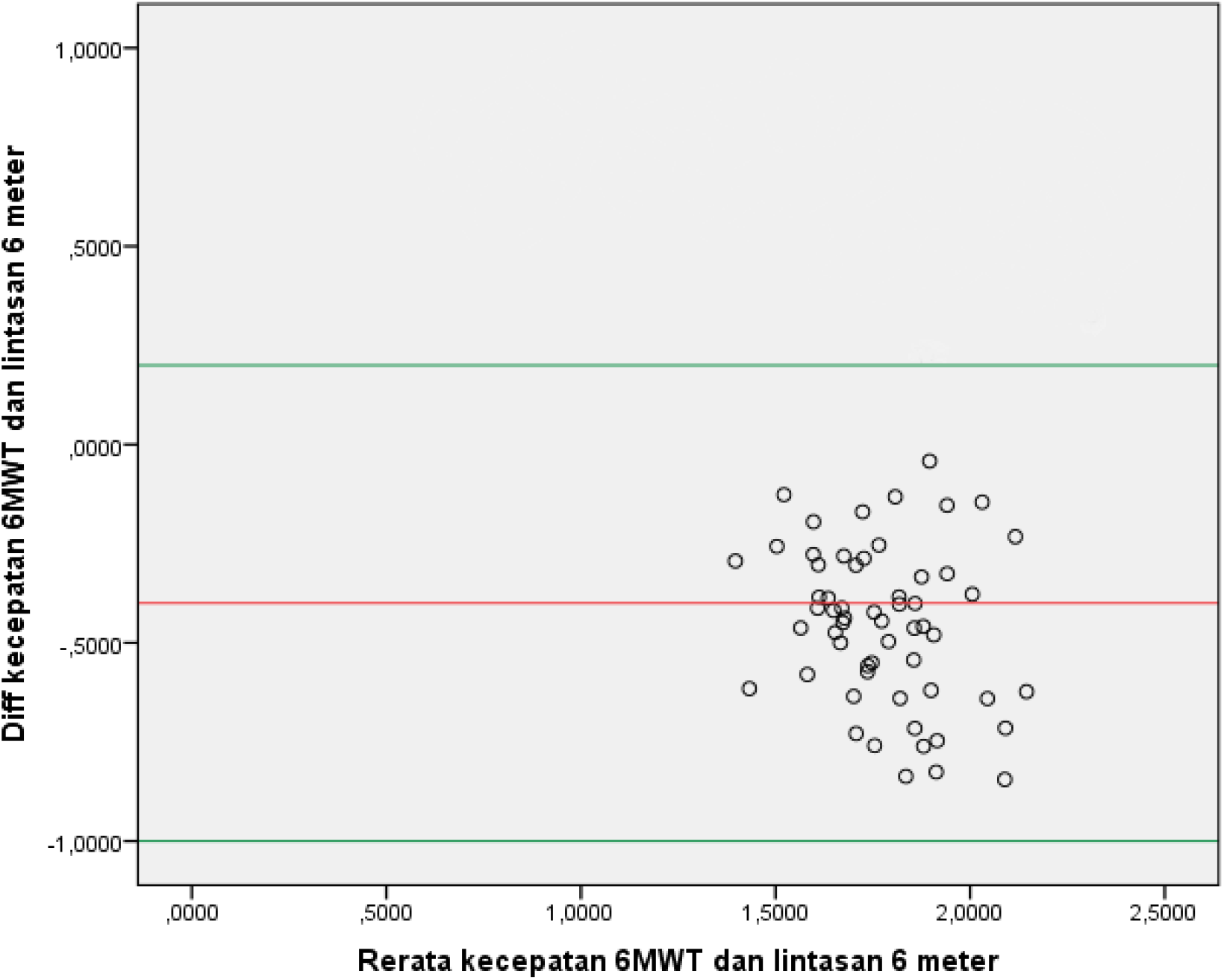
Uji Kesesuaian Bland-Altman kecepatan berjalan pada Uji Jalan Enam Menit dengan Uji Jalan Empat Meter, pada Lintasan Enam Meter

**Grafik 2.**
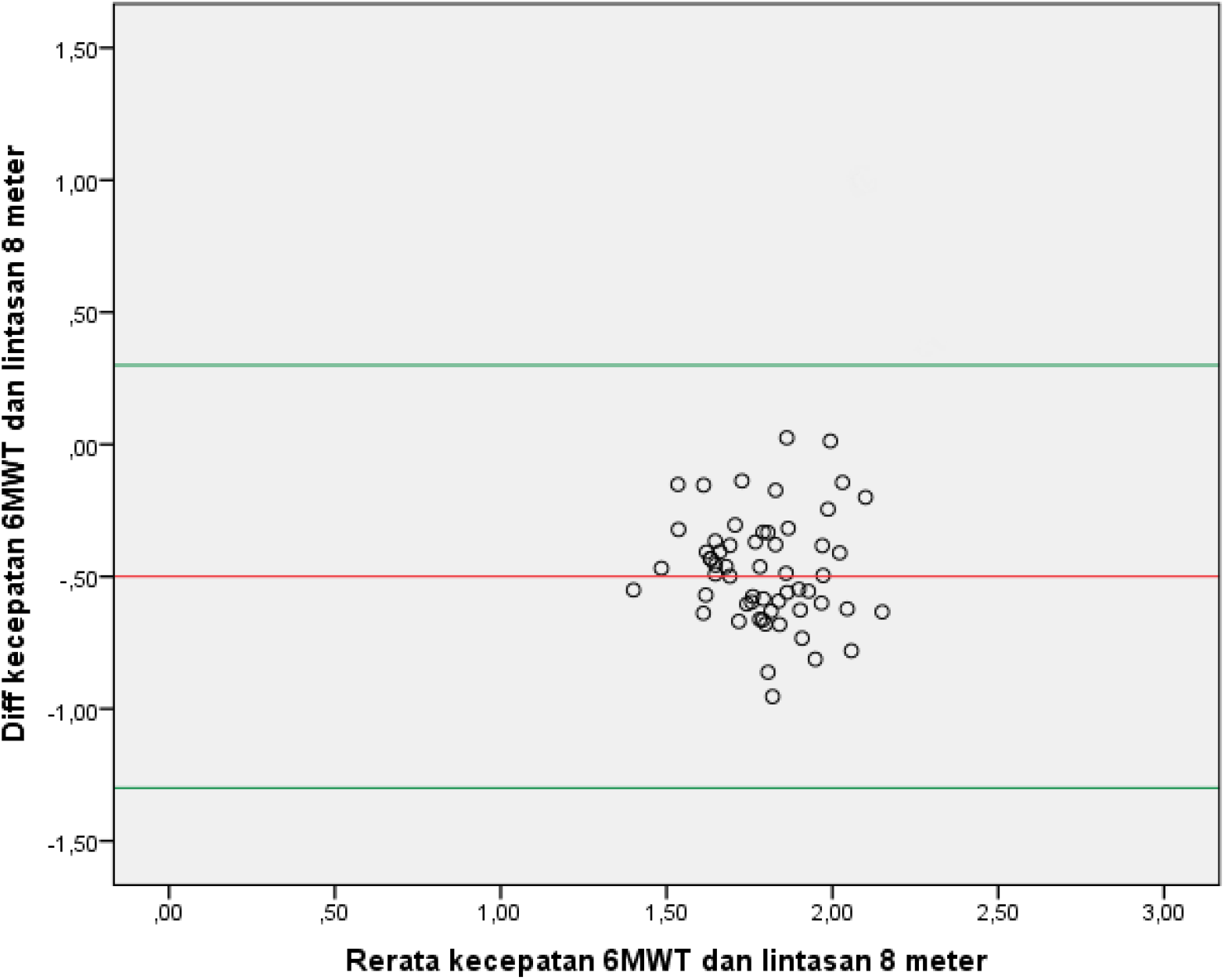
Uji Kesesuaian Bland-Altman Uji kecepatan berjalan Jalan Enam Menit dengan Uji Jalan Empat Meter, pada Lintasan Delapan Meter

